# StainAI-Web: An Interactive Platform for Whole-Slide Microglial Morphology Analysis

**DOI:** 10.64898/2026.10.17.759834

**Authors:** Da-Yuan Liu, Hsiang-Jou Chuang, Chao-Hsiung Hsu, Be-Ming Chang, EssietAdid Ette, Mujan Shih, Yueh-An Liao, Hania Zehra, Sakinah Ali, Tsang-Rung Tu, Stephen Lin, Paul C. Wang, Michael Shoykhet, Tsang-Wei Tu

## Abstract

Microglial morphology analysis has emerged as a critical approach in neuroinflammation research. The increasing availability of high-resolution and whole-slide histology images has created strong demand for robust, scalable analytical methods capable of efficiently processing these datasets. Existing approaches may require code-based local computation or prolonged processing times for whole-slide images, while manual morphology quantification remains labor-intensive. To address these limitations, we introduce StainAI-Web, a server that performs rapid, automated microglial morphology detection across whole-slide immunohistochemistry images in the cloud. The system provides interactive grayscale preprocessing and supports ROI-based regional quantification, allowing users to define specific tissue regions and systematically compare microglial populations across regions. An interactive visualization module dynamically updates quantitative summaries of morphological distributions as ROIs are selected or modified. Analysis results, including structured JSON files and morphology-annotated overlays, are downloadable for downstream analysis or integration into external computational pipelines. These integrated features provide an accessible framework for efficient and reproducible whole-slide microglial morphology analysis. The StainAI-Web is available at https://stainaiviewer.azurewebsites.net/.

## Introduction

Microglia exhibit a range of morphologies, from highly ramified cells with small somata and extensive processes to hypertrophic, bushy, amoeboid, and dystrophic forms(1–3). These morphological patterns vary with local environment, brain region, injury or disease stage, and inflammatory conditions, making morphology a useful dimension for studying microglial responses and neuroinflammation(4, 5). Quantifying this diversity at the cellular level, however, requires analysis that can preserve both the morphology of individual cells and their spatial organization across tissue. The scale of modern histological imaging has also made this challenge increasingly important. Whole-slide imaging generates high-resolution images spanning large tissue areas and containing thousands to millions of microglial cells. Yet, existing approaches remain limited in scale: semi-manual analysis becomes increasingly laborious and difficult to reproduce, while field-of-view-based automated tools may overlook regional heterogeneity and introduce sampling bias, limitations that become increasingly consequential as image size and cohort size expand(6–9). The challenge therefore extends beyond cell counting to transforming large-scale histological images into quantitative cellular and spatial information that can be readily interrogated across tissue(10, 11).

Deep learning has substantially advanced automated microglial detection, segmentation, and morphology classification(12, 13). We previously introduced StainAI, a deep-learning framework that integrates cell detection, segmentation, morphology classification, and atlas-based brain mapping for automated whole-slide analysis of Iba1-stained microglia. StainAI enables high-throughput characterization of microglia from low-magnification (∼20X) whole-brain images while preserving spatial context, revealing both established and novel activation patterns following cardiac arrest-induced brain injury. However, adoption of sophisticated image-analysis pipelines remains challenging outside computationally specialized laboratories, as approaches often require local software installation, programming expertise, dedicated hardware, parameter configuration, and manual data processing(14). Researchers may also need to integrate multiple computational and visualization tools to obtain quantitative results, creating additional barriers between image acquisition and biological interpretation.

To address these challenges, we developed StainAI-Web, an interactive web-based platform built on the StainAI framework that brings automated whole-slide image analysis into an integrated browser-based workflow(12). The platform incorporates adaptive image preprocessing, optimized large-image handling, and overlap-aware patch-based inference to enable efficient analysis of high-resolution histological images through a globally accessible web interface(12). This study aims to establish a scalable framework for whole-slide microglial analysis that reduces computational barriers and streamlines quantitative morphological interpretation.

## Materials and methods

### System Back-end Framework

StainAI-Web was designed as a modular framework integrating large-scale image processing, deep learning detection, and web-based visualization and quantification into a unified pipeline (Fig. 1). In Stage 1, JPEG or PNG images were prepared for detection using user-specified spatial resolution and grayscale preprocessing parameters. When spatial resolution was supplied in µm/pixel, image dimensions were scaled by the ratio of the supplied resolution to the model’s training resolution of 0.464 µm/pixel. A supplied resolution of 0.464 µm/pixel resulted in no change in spatial dimensions. When resolution was omitted, the original image dimensions were retained, with 0.464 µm/pixel assumed for inference only.

**Figure 1.**
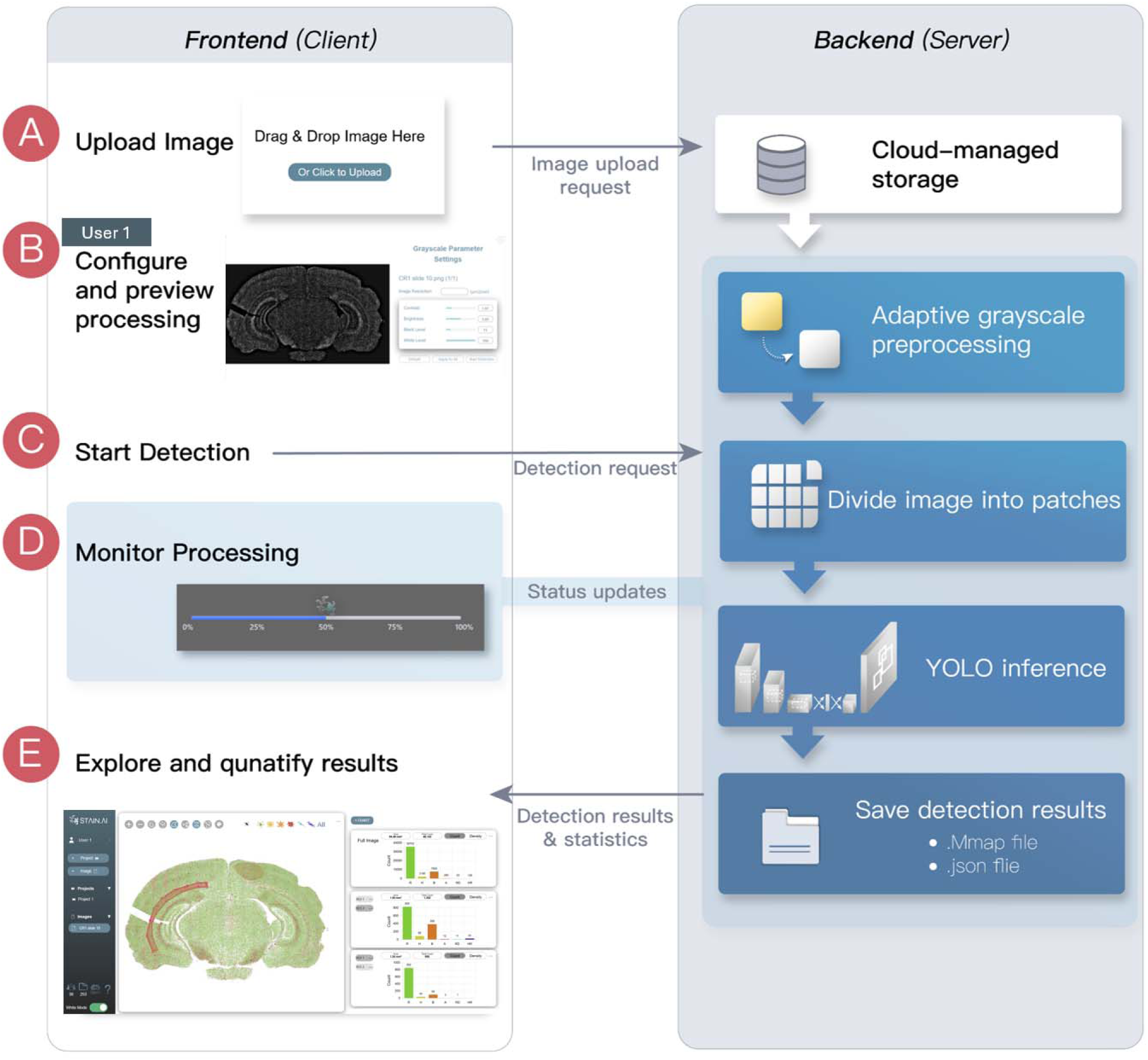
The StainAI-Web workflow integrating browser-based analysis with server-side processing. (A) Users upload microscopy images to cloud-managed storage. (B) Preprocessing parameters are adjusted and previewed before analysis. (C) The backend performs adaptive grayscale preprocessing, patch image processing, and YOLO-based inference. (D) Processing status is communicated to the frontend in real time. (E) Detection results and summary statistics are returned for interactive visualization and quantification, while output data are stored as .mmap and .json files for subsequent access.

The detection image subsequently underwent modality-specific grayscale preprocessing, with the processing pathway automatically selected based on image border intensity characteristics. Brightfield RGB images were converted to luminance and inverted, whereas an optional fluorescence pathway performed channel selection based on signal-to-background characteristics. Both pathways included large-radius local background correction to reduce slowly varying background intensity variations(15), followed by percentile normalization, gamma correction(16), and gain adjustment. User-specified lower and upper percentiles defined the intensity range mapped to the normalized scale, gamma controlled nonlinear intensity mapping, and gain multiplicatively scaled the resulting intensities.

In Stage 2, the processed grayscale images were analyzed using an overlap-aware, patch-based detection pipeline optimized for ultra-high-resolution whole-slide images(10, 12, 17). Each image was divided into 640 × 640-pixel patches, with coordinates generated along both spatial dimensions using a predefined stride of:

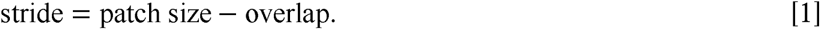

The overlap was set to 320 pixels, corresponding to half of the patch size, to provide redundant coverage near patch boundaries and ensure that cells near the edges could be captured in adjacent patches(12). Patch starting positions were generated on a regular grid, with a terminal position added when necessary for complete image coverage. For dimensions smaller than the patch size, a single patch was generated along that axis. Boundary patches were adjusted to remain within valid image bounds or padded to the target size when required. Overlapping patches reduced boundary-related artifacts by improving capture of edge-located cells and facilitating consolidation of redundant detections. Patch-based processing avoided downsampling the entire whole-slide image to a single YOLO input, thereby preserving cellular morphology and spatial detail while maintaining compatibility with the detection model(18).

The original StainAI pipeline sequentially combined YOLO-based detection(18), UNet segmentation(19), morphometric feature extraction(12), and decision-tree classification(20). Although this hybrid approach enabled detailed single-cell segmentation and morphometric analysis, its multiple processing stages increased computational complexity. StainAI-Web instead used a YOLO-only model to directly localize microglia and classify them into one of six morphological classes: ramified (R), hypertrophic (H), bushy (B), amoeboid (A), rod-shaped (RD), and hypertrophic rod-shaped (HR). This simplified architecture reduced inference complexity while supporting efficient whole-slide processing on a shared web server (12, 21).

The StainAI-Web model was initialized from pretrained YOLO12x weights and fine-tuned on annotated Iba1-stained whole-brain images from the original StainAI dataset, including images from control rats without cardiac arrest (Control) and rats subjected to 12-min asphyxial cardiac arrest followed by resuscitation (CA12)(12). Training bounding boxes were derived from segmentation masks originally generated for training StainAI, while morphology labels were assigned by the StainAI C5.0 classifier.

Source images were approximately 30,000 × 20,000 pixels at 0.464 µm/pixel and were divided into 640 × 640-pixel patches. Patches underwent horizontal flipping, rotations of 30° − 270°, translation, shear, and mosaic augmentation and were randomly divided into training and held-out evaluation sets at an 80:20 ratio. The model was trained for 300 epochs with a batch size of 12, automatic optimizer selection, a three-epoch warm-up, and a fixed random seed of 0. Checkpoints were saved every 50 epochs, with training and validation curves recorded for performance monitoring.

During inference, overlapping patches were processed in batches, with automatic batch-size reduction in response to GPU memory limitations. Retained bounding boxes were mapped to whole-slide coordinates and clipped to their corresponding patch boundaries. Predictions were consolidated using class-specific non-maximum suppression, followed by class-independent IoU clustering and confidence-weighted box fusion(22). Each cluster yielded a representative bounding box assigned to the morphology class with the highest summed confidence, reducing redundant detections and conflicting class assignments across overlapping patches.

In Stage 3, aggregated detections were transformed into structured outputs for quantitative analysis and visualization. Brenner’s focus measure (FM) was calculated from pixel intensities within each bounding box to quantify local image sharpness(23), while a microglial activation score (MAS) was calculated using morphology-specific weights(12):

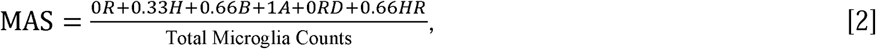

Where *R*,*H*, *B*, *A*, *RD*, and *HR* denote the counts of ramified, hypertrophic, bushy, ameboid, rod-shaped, and hypertrophic rod-shaped microglia, respectively. The weighting coefficients reflected increasing levels of microglial activation across morphological states(12). The resulting MAS was bounded between 0 and 1 by construction, enabling quantitative comparison across ROIs and samples(12).

Detection results were consolidated into a downloadable JSON file containing each detection’s bounding box coordinates, centroid location, morphology class, and focus measure (FM), together with analysis-level microglial activation scores (MAS), supporting downstream analysis and reuse. Visualization outputs overlaid semi-transparent, class-specific regions on the original and grayscale images using alpha compositing rather than conventional bounding boxes. Full-slide morphological statistics were aggregated across classes and displayed as a summary bar chart. In addition to the default global quantification,

ROI-based analysis was also supported, with charts generated from user-defined selections while retaining the full-image chart as a global reference. Two quantitative modes were provided: Count, representing raw cell counts by morphology class, and Density, representing cell counts normalized by the analyzed ROI area. The ROI area was calculated as:

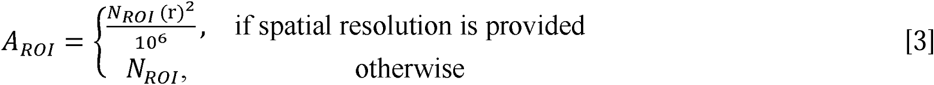

Here,*N_roi_* denotes the number of pixels within the ROI at the original uploaded image scale, and r is the user-supplied spatial resolution in µm/pixel. Accordingly,*A_roi_* was expressed in mm² when spatial resolution was supplied and in px² otherwise. Density was calculated by dividing the cell count for each morphology class by *A_roi_* and was expressed in cells/mm² or cells/px², respectively. The default resolution assumed for inference was not used for physical area conversion.

Following analysis, four downloadable outputs were generated: a morphology-class count chart (.png); a preprocessing record containing grayscale settings, spatial resolution, and image area (.txt); structured results containing analysis metadata and per-cell coordinates, confidence scores, morphology classes, focus measures, and MAS values (.json); and a two-page TIFF containing the original image and its color-coded morphology overlay (.tif). Outputs were organized by source image in cloud storage for retrieval and downstream analysis.

### System Front-end Interface

StainAI-Web provided a browser-based interface for microglial morphology analysis without local installation. The workflow comprised two sequential views: an upload/configuration view and an interactive analysis viewer. Users uploaded whole-slide images and adjusted grayscale preprocessing parameters, including contrast, brightness, and black/white levels, with real-time preview. When image resolution metadata (µm/pixel) were provided by the user, the system supported resampling toward the model training scale of 0.464 µm/pixel. Confirmed settings were then transmitted to the backend for preprocessing and detection, with progress displayed before automatic transition to the analysis viewer.

The analysis viewer comprised a project and image management panel, a central image canvas, and a quantitative visualization panel (Fig. 2). The Projects and Images sections allowed users to reloaded analysis and organize images into custom folders by experiment, subject, or tissue category, with options to create, move, rename, delete, and drag and drop items. The OpenSeadragon-based canvas (24) supported panning and zooming of high-resolution images and displayed semi-transparent overlays for six predicted morphologies. An all-cell display mode showed all detections with a uniform overlay while retaining morphology-resolved quantitation.

**Figure 2.**
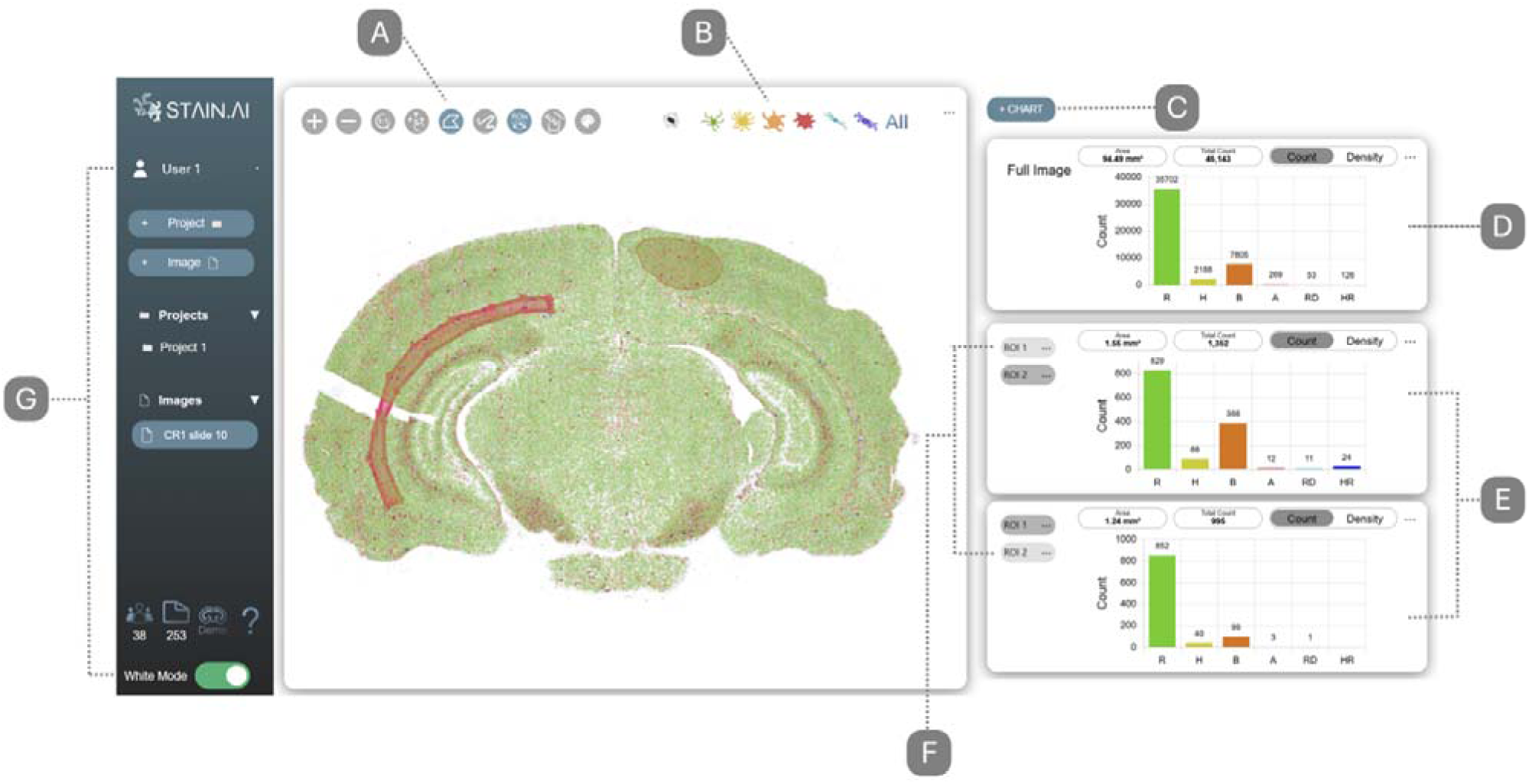
Interactive visualization and analysis interface of StainAI-Web. (A) Image navigation and ROI toolbar supports zooming, panning, drawing, editing, and displaying regions of interest. (B) Morphology display controls enable selectively viewing of individual classes, all morphologies, or all detected cells. (C) Additional ROI-linked quantification panels can be created using the chart control. (D) Whole-slide quantification summarizes morphology distributions, analyzed area, total cell count, and count- or density-based measurements. (E) ROI panels enable side-by-side quantitative comparison of selected regions. (F) ROI management panel allows regions to be selected, renamed, and deleted. (G) Project and image management panel organizes uploaded images and analysis results.

A Konva-based ROI layer (25) enabled synchronized ROI drawing, editing and navigation. Slide-level detections were rescaled to display coordinates and synchronized with ROI selections and morphology filters. The visualization panel provided whole-slide and ROI-based Count and Density measurements, including analyzed area and total cell counts. ROIs could be named, selected, recolored, edited, deleted, or combined for regional analysis. Quantification was based on detected cell centers and updated dynamically as ROIs were modified. Results and ROI information could be exported for reporting and multi-slide analysis

### Application and Performance Evaluation

Representative rat whole-brain Iba1 immunohistochemistry images from previously published control and cardiac arrest datasets(12, 26). Cardiac arrest provided a biologically relevant use case for evaluating whole-slide morphology detection, quantitative analysis, and ROI-based regional assessment of microglial activation (26, 27). Images were processed through the complete StainAI-Web workflow, including preprocessing, patch-based detection, morphology classification, whole-slide quantification, and interactive ROI analysis.

Model performance was evaluated on an independent held-out evaluation patches from the two-source whole-slide images using precision, recall, F1 score, mAP@0.5, and mAP@0.5:0.95, overall and by morphology class. Total microglial counts were compared between StainAI-Web and the original StainAI framework across five representative whole-slide images, while morphology distributions were compared for CR1 Slide 10. Relative differences in total counts and percentage-point differences in class-specific proportions were calculated. Computational scalability was evaluated using five whole-brain images ranging from 19.2 to 438.9 megapixels, randomly selected from one control (CR1) and three cardiac arrest brains (N25, N31, N13), and processing time was recorded for each image. Finally, regional analysis used five user-defined ROIs encompassing the left and right external capsules, corpus callosum, and bilateral cortices to demonstrate regional comparisons of microglial morphology in representative control and cardiac arrest images.

## Results

### Use Case Demonstration

StainAI-Web generated morphology-resolved maps across entire whole-brain sections, allowing microglial morphotypes to be visualized in their original anatomical context. The control section (27,160 × 16,160 pixels; 438.9 megapixels) contained 46,143 detected microglia, compared with 40,032 in the cardiac arrest section (25,802 × 15,150 pixels; 390.9 megapixels). Processing required 15 min 30 s and 12 min 50 s, respectively, on an Azure Premium P3v3 host. In these representative sections, whole-brain analysis showed a marked difference in morphology distributions between the control and cardiac arrest images. The proportion of ramified cells was 77.4% in the control section and 33.0% in the cardiac arrest section, whereas non-ramified morphotypes accounted for 22.6% to 67.0%, respectively. Bushy microglia showed the largest difference in proportion, accounting for 7,805 (16.9%) of detected cells in the control section and 23,019 (57.5%) in the cardiac arrest section. The corresponding counts and proportions were 269 (0.6%) to 1,509 (3.8%) for amoeboid microglia, and 126 (0.3%) and 443 (1.1%) for hypertrophic rod-shaped microglia.

### Model Performance and Comparison with StainAI

Evaluation on held-out patches from the two-source whole-slide images showed high detection performance, with overall precision, recall, and F1 scores of 0.970, 0.955, and 0.962, respectively. mAP@0.5 and mAP@0.5:0.95 were 0.975 and 0.904 (Table 1). Across the six morphology classes, F1 scores ranged from 0.945 for hypertrophic to 0.982 for hypertrophic rod-shaped microglia; precision exceeded 0.95 for all classes, and mAP@0.5 ranged from 0.964 to 0.985. Across five representative whole-slide images, StainAI-Web produced broadly comparable total microglial counts to the original StainAI framework (Table 2). The mean relative difference was 9.66%, ranging from 7.08% for N13 Slide 1 to 14.23% for CR1 Slide 2. For CR1 Slide 10, StainAI detected 42,439 microglia, whereas StainAI-Web detected 46,143 cells, an 8.73% relative difference. Morphology distributions showed greater differences between the two methods. In CR1 Slide 10, ramified cells accounted for 58.7% with StainAI and 77.3% with StainAI-Web, while hypertrophic cells accounted for 14.0% and 4.7%, and bushy cells for 9.9% and 16.9%, respectively. Amoeboid and rod-shaped cells differed by 8.1 and 8.0 percentage points, respectively, whereas hypertrophic rod-shaped cells differed by only 0.2 percentage points.

**Table 1.**
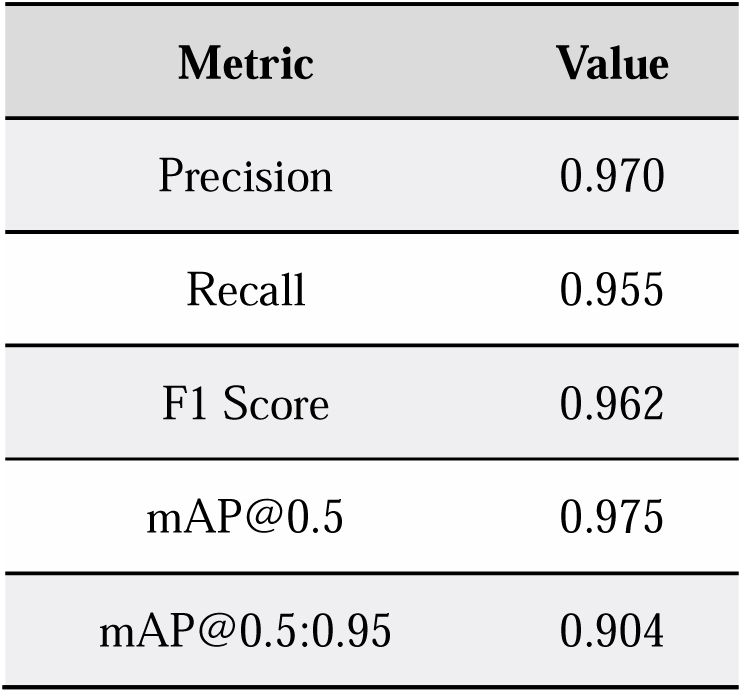
Overall detection performance of StainAI-Web on an independent test dataset. Precision, recall, F1 score, mAP@0.5, and mAP@0.5:0.95 summarize the detection performance across all six microglial morphology classes.

**Table 2.** Benchmark comparison of total microglial counts between StainAI-Web and the original StainAI framework. Total microglial counts, absolute differences, and relative differences were compared across five representative whole-slide images, with mean values reported across images.

| Image | StainAI, n | StainAI-Web, n | Difference, n | Relative difference (%) |
| --- | --- | --- | --- | --- |
| CR1 Slide 10 | 42439 | 46143 | +3704 | 8.73 |
| N25 Slide 10 | 43355 | 40032 | -3323 | 7.66 |
| CR1 Slide 2 | 22119 | 25267 | +3148 | 14.23 |
| N31 Slide 3 | 14325 | 15841 | +1516 | 10.58 |
| N13 Slide 1 | 2273 | 2434 | +161 | 7.08 |
| Mean | 24902.2 | 25943.4 | +1041.2 | 9.66 |

### Computational Scalability

Processing time increased with whole-slide image size across five images ranging from 19.2 to 438.9 megapixels (Table 3). Runtime ranged from 32 s for the smallest image (19.2 megapixels; 3,628 × 5,281 pixels) to 15 min 30 s for the largest image (438.9 megapixels; 27,160 × 16,160 pixels). Images of 131.7, 214.0, and 390.9 megapixels required 3 min 30 s, 5 min 59 s, and 12 min 50 s, respectively. Processing rates remained within 1.59–2.12 s per megapixel, indicating predictable scaling across image sizes.

**Table 3.** Computational scalability of StainAI-Web across whole-slide images of varying sizes. Image dimensions, file sizes, and processing times for five representative images demonstrate runtime scaling with image size.

| Image | Dimensions (pixels) | Image size (mb) | Total runtime (min:sec) |
| --- | --- | --- | --- |
| CR1 Slide 10 | 27160 x 16160 | 516 | 15:30 |
| N25 Slide 10 | 25802 x 15150 | 417 | 12:50 |
| CR1 Slide 2 | 17654 x 12120 | 35.6 | 05:59 |
| N31 Slide 3 | 10864 x 12120 | 20.6 | 03:30 |
| N13 Slide 1 | 3628 x 5281 | 5 | 00:32 |

### ROI-Based Regional Analysis

ROI analysis of representative control and cardiac arrest sections showed differences in microglial morphology distributions across five selected regions. In all five regions analyzed, the proportion of ramified microglia was lower in the cardiac arrest section, whereas activated morphotypes morphotypes accounted for a larger proportion of detected cells. The corpus callosum showed the largest difference, with activated morphotypes accounting for 19.91% of detected cells in the control section and 91.56% in the cardiac arrest section. Both left and right cortices showed similar difference, from approximately 15.6% in the control image to approximately 77.0% in the cardiac arrest image. Area-normalized analysis reported higher activated microglial densities in the cardiac arrest section than in the control section: left external capsule (8,994 to 14,731 cells/mm²), right external capsule (10,900 to 15,131 cells/mm²), corpus callosum (2,684 to 13,238 cells/mm²), and bilateral cortices (approximately 2,100–2,200 to 8,100–8,500 cells/mm²). These results demonstrate that StainAI-Web supports both whole-slide and region-specific quantitative analysis of microglial morphology.

## Discussion

### An Accessible Platform for Whole-Slide Microglial Analysis

StainAI-Web provides an accessible web-based workflow for automated whole-slide microglial morphology analysis. To our knowledge, it is the first platform to integrate whole-slide inference, morphology classification, interactive visualization, ROI-based quantification, and structured data export within a free, unified environment. This integration addresses limitations of workflows that rely on manual analysis, local ImageJ/Fiji pipelines, or custom computational tools, which can limit accessibility, scalability, and reproducibility(3, 4, 6–9, 14). A key advancement of StainAI-Web is the redesign of the original StainAI computational pipeline for web-server deployment. StainAI uses a multi-stage architecture, combining YOLO-based detection, UNet segmentation, morphometric feature extraction, and decision-tree classification(12). StainAI-Web instead uses a streamlined YOLO-based model that directly detects and classifies six microglial morphologies using overlap-aware patch processing for large whole-slide images.

This simplifies computational workflow while retaining the six morphological categories used in StainAI. Thus, the two plarforms serve complementary purposes: StainAI provides detailed segmentation, morphometric analysis, and atlas-based mapping, whereas StainAI-Web prioritizes accessible and efficient whole-slide analysis. By integrating preprocessing, inference, visualization, and result generation through the browser and server, StainAI-Web reduces the need for specialized software, dedicated GPUs, and deep-learning expertise(12, 28).

### Architectural Trade-offs and Computational Performance

StainAI-Web produced broadly comparable total microglial counts to the original StainAI framework (Table 2), but differences were observed in their morphology distributions. These differences may partly reflect the distinct computational architectures of the two systems. The original StainAI pipeline integrates multiple stages of detection, segmentation, morphometric feature extraction, and decision-tree classification, where errors introduced at individual stages may propagate through subsequent processing steps. Furthermore, feature selection and classification methods can influence the final morphology assignments, making it difficult to assess the contributions of individual processing stages to classification uncertainty. In contrast, StainAI-Web directly predicts microglial locations and morphology classes using a single YOLO model, reducing dependencies between processing stages and simplifying computational deployment. The YOLO-based model also provides confidence scores associated with individual detections and their predicted morphology classes, offering additional information for assessing prediction reliability and identifying low-confidence detections for further review. These architectural differences highlight the trade-offs between detailed morphometric characterization and streamlined morphology classification. By eliminating separate segmentation and morphometric feature-extraction stages, the YOLO-only architecture reduces computational overhead and enables rapid whole-slide processing on a shared server, with whole-slide processing times reported in Table 3.

### Interactive Whole-Slide Visualization and ROI-Based Analysis

The control-versus-cardiac-arrest application demonstrated the utility of combining whole-slide and ROI-based analysis. The observed differences in microglial morphology distributions between representative control and cardiac arrest sections were broadly consistent with previously reported findings (13, 29, 30).

The interactive viewer allows users to visually compare the original histological image with morphology-classified overlays when defining and adjusting ROIs, facilitating the identification of anatomical regions and local morphological patterns. Because ROIs are analyzed after whole-slide inference, regional cell counts and morphology distributions can be updated dynamically as ROIs are modified, without rerunning detection. (12, 27, 28) Combining cell counts with area-normalized densities also facilitates comparisons among regions of different sizes. This integration of whole-slide visualization and flexible regional quantification provides a practical approach for investigating spatial heterogeneity in microglial responses across anatomical and pathological conditions(29, 30).

### Image Quality and Preprocessing for Reliable Morphology Analysis

IHC image appearance can vary substantially with antibody characteristics, staining conditions, camera sensitivity, exposure, and post-processing, creating an important source of variability for deep-learning–based analysis(10). Because detection and morphology classification are sensitive to image contrast and brightness, inconsistent image quality or inappropriate preprocessing may affect classification. StainAI-Web addresses this variability through modality-aware grayscale conversion, background correction, normalization, and interactive contrast adjustment, allowing users to preview and optimize preprocessing before inference. Image quality remains an important factor in reliable morphology analysis, as preprocessing cannot fully recover cellular details lost due to poor focus, insufficient spatial resolution, or signal saturation. In addition to the built-in preprocessing tools provided by StainAI-Web, users may preprocess images using ImageJ/Fiji or image-processing software supplied by microscope manufacturers before uploading them to the server. These controls require careful use. Excessive background suppression or contrast enhancement may remove fine cellular processes or alter morphological features, potentially affecting detection and classification results. Preprocessing settings should also be applied consistently across samples intended for comparison to minimize systematic variability. Standardized outputs, including morphology maps, structured JSON files, quantitative summaries, and preprocessing parameters, further support reproducible downstream analysis.

### Limitations

StainAI-Web is currently limited to microglial morphology analysis and requires additional training for other cell types. Its streamlined YOLO-only architecture prioritizes deployment efficiency and therefore does not provide the detailed segmentation, morphometric measurements, atlas registration, or three-dimensional mapping available in the original StainAI framework(12). Differences in morphology distributions between StainAI and StainAI-Web also indicate that class-specific results from the two pipelines may not be directly interchangeable. Broader validation across independent datasets, imaging systems, staining protocols, and disease models is also needed, as variations in image quality and resolution may affect performance.

Although StainAI-Web supports fluorescence image preprocessing, detection and classification performance on fluorescence images remains to be validated. The current model evaluation was primarily based on patches derived from two source whole-slide images. Potential overlap of augmented cellular content between training and evaluation sets may lead to overestimated performance metrics, highlighting the need for independent validation. The control-versus-cardiac-arrest comparisons were based on representative sections and should therefore be interpreted as demonstrations of the analytical workflow rather than group-level biological findings. Processing time will vary with image size and server resources. Finally, morphology-based classification represents a structural indicator of microglial state rather than a direct measure of molecular or inflammatory function(31).

### Future Development

Future work will expand StainAI-Web to additional cell types, improve model robustness, and strengthen external validation. Selective integration of segmentation and morphometric analysis could extend its analytical capabilities while balancing additional computational demands with web-server efficiency(12). Further development will address heterogeneous imaging conditions, expand quantitative measurements, improve ROI definition and comparison across images, and benchmark computational performance across image sizes, server configurations, and user loads. Model version tracking may also be incorporated to support reproducibility and longitudinal consistency as detection models are updated.

## Author contributions

Da-Yuan Liu (Methodology [lead], Software [lead], Formal analysis [lead], Validation [lead], Visualization [lead], Writing – original draft [lead]), Hsiang-Jou Chuang, (Methodology [equal], Data curation [equal], Validation [equal], Writing – review [equal]), Chao-Hsiung Hsu, (Methodology [equal], Data curation [equal], Writing – review [equal]), Be-Ming Chang, (Conceptualization [equal], Methodology [equal], Data curation [equal], Writing – review [equal]), EssietAdid Ette, (Methodology [equal], Data curation [equal], Writing – review [equal]), Mujan Shih, (Methodology [equal], Data curation [equal], Writing – review [equal]), Yueh-An Liao, (Methodology [equal], Data curation [equal], Writing – review [equal]), Hania Zehra, (Investigation [supporting], Validation [supporting], Writing – review [equal]), Sakinah Ali, (Investigation [supporting], Validation [supporting], Writing – review [equal]), Tsang-Rung Tu, (Investigation [supporting], Validation [supporting], Writing – review [equal]), Stephen Lin, (Project administration [supporting], Writing – review [equal]), Paul C. Wang, (Project administration [equal], Funding acquisition [equal], Resources [equal], Writing – review [equal]), Michael Shoykhet, (Conceptualization [equal], Funding acquisition [equal], Resources [equal], Writing – review & editing [equal]), Tsang-Wei Tu (Conceptualization [lead], Supervision [lead], Project administration [lead], Funding acquisition [lead], Resources [equal], Methodology [equal], Validation [equal]), Writing – review & editing [lead])

## Data Availability

StainAI-Web is freely accessible for non-commercial academic and research purposes at https://stainaiviewer.azurewebsites.net/.

## Reference

1. Savage, J.C., Carrier, M. and Tremblay, M.È. (2019) Morphology of Microglia Across Contexts of Health and Disease. In Methods in Molecular Biology.Vol. 2034. 10.1007/978-1-4939-9658-2_2

2. Colonna, M. and Butovsky, O. (2017) Microglia function in the central nervous system during health and neurodegeneration. Annu. Rev. Immunol., 35. 10.1146/annurev-immunol-051116-052358

3. Green, T.R.F. and Rowe, R.K. (2024) Quantifying microglial morphology: an insight into function. Clin. Exp. Immunol., 216. 10.1093/cei/uxae023

4. Reddaway, J., Richardson, P.E., Bevan, R.J., Stoneman, J. and Palombo, M. (2023) Microglial morphometric analysis: so many options, so little consistency. Front. Neuroinform., 17. 10.3389/fninf.2023.1211188

5. Vidal-Itriago, A., Radford, R.A.W., Aramideh, J.A., Maurel, C., Scherer, N.M., Don, E.K., Lee, A., Chung, R.S., Graeber, M.B. and Morsch, M. (2022) Microglia morphophysiological diversity and its implications for the CNS. Front. Immunol., 13. 10.3389/fimmu.2022.997786

6. Green, T.R.F., Murphy, S.M. and Rowe, R.K. (2022) Comparisons of quantitative approaches for assessing microglial morphology reveal inconsistencies, ecological fallacy, and a need for standardization. Sci. Rep., 12. 10.1038/s41598-022-23091-2

7. Leyh, J., Paeschke, S., Mages, B., Michalski, D., Nowicki, M., Bechmann, I. and Winter, K. (2021) Classification of Microglial Morphological Phenotypes Using Machine Learning. Front. Cell. Neurosci., 15. 10.3389/fncel.2021.701673

8. Clarke, D., Crombag, H.S. and Hall, C.N. (2021) An open-source pipeline for analysing changes in microglial morphology. Open Biol., 11. 10.1098/rsob.210045

9. Stetzik, L., Mercado, G., Smith, L., George, S., Quansah, E., Luda, K., Schulz, E., Meyerdirk, L., Lindquist, A., Bergsma, A., et al. (2022) A novel automated morphological analysis of Iba1+ microglia using a deep learning assisted model. Front. Cell. Neurosci., 16. 10.3389/fncel.2022.944875

10. Dimitriou, N., Arandjelović, O. and Caie, P.D. (2019) Deep Learning for Whole Slide Image Analysis: An Overview. Front. Med. (Lausanne)., 6. 10.3389/fmed.2019.00264

11. Kim, J., Pavlidis, P. and Ciernia, A.V. (2024) Development of a High-Throughput Pipeline to Characterize Microglia Morphological States at a Single-Cell Resolution. eNeuro, 11. 10.1523/ENEURO.0014-24.2024

12. Hsu, C.H., Hsu, Y.Y., Chang, B.M., Raffensperger, K., Kadden, M., Ton, H.T., Ette, E.A., Lin, S., Brooks, J., Burke, M.W., et al. (2025) StainAI: quantitative mapping of stained microglia and insights into brain-wide neuroinflammation and therapeutic effects in cardiac arrest. *Commun*. Biol., 8. 10.1038/s42003-025-07926-y http://www.ncbi.nlm.nih.gov/pubmed/40114030

13. Stetzik, L., Mercado, G., Smith, L., George, S., Quansah, E., Luda, K., Schulz, E., Meyerdirk, L., Lindquist, A., Bergsma, A., et al. (2022) A novel automated morphological analysis of Iba1+ microglia using a deep learning assisted model. Front. Cell. Neurosci., 16. 10.3389/fncel.2022.944875

14. Aeffner, F., Zarella, M.D., Buchbinder, N., Bui, M.M., Goodman, M.R., Hartman, D.J., Lujan, G.M., Molani, M.A., Parwani, A. V., Lillard, K., et al. (2019) Introduction to digital image analysis in whole-slide imaging: A white paper from the digital pathology association. J. Pathol. Inform., 10. 10.4103/jpi.jpi_82_18

15. Sternberg, S.R. Biomedical Image Processing.

16. Johnson, J. (2012) Not seeing is not believing: improving the visibility of your fluorescence images. Mol. Biol. Cell, 23, 754–757. 10.1091/MBC.E11-09-0824 http://www.ncbi.nlm.nih.gov/pubmed/22379119

17. Berman, A.G., Orchard, W.R., Gehrung, M. and Markowetz, F. (2023) SliDL: A toolbox for processing whole-slide images in deep learning. PLoS One, 18. 10.1371/journal.pone.0289499

18. Redmon, J., Divvala, S., Girshick, R. and Farhadi, A. (2016) You only look once: Unified, real-time object detection. In Proceedings of the IEEE Computer Society Conference on Computer Vision and Pattern Recognition.Vol. 2016-December. 10.1109/CVPR.2016.91

19. Falk, T., Mai, D., Bensch, R., Çiçek, Ö., Abdulkadir, A., Marrakchi, Y., Böhm, A., Deubner, J., Jäckel, Z., Seiwald, K., et al. (2019) U-Net: deep learning for cell counting, detection, and morphometry. Nat. Methods, 16. 10.1038/s41592-018-0261-2

20. Max Kuhn, M. (2026) Package ‘C50’ Type Package Title C5.0 Decision Trees and Rule-Based Models.

21. Redmon, J., Divvala, S., Girshick, R. and Farhadi, A. (2016) You only look once: Unified, real-time object detection. In Proceedings of the IEEE Computer Society Conference on Computer Vision and Pattern Recognition. IEEE Computer Society, Vol. 2016-December, pp. 779–788. 10.1109/CVPR.2016.91

22. Solovyev, R., Wang, W. and Gabruseva, T. (2021) Weighted boxes fusion: Ensembling boxes from different object detection models. Image Vis. Comput., 107, 104117. 10.1016/J.IMAVIS.2021.104117

23. Brenner, J.F., Dew, B.S., Horton, J.B., King, T., Neurath, P.W. and Selles, W.D. (1976) An automated microscope for cytologic research: a preliminary evaluation. Journal of Histochemistry and Cytochemistry, 24. 10.1177/24.1.1254907

24. OpenSeadragon.

25. Konva - JavaScript Canvas 2d Library.

26. Ton, H.T., Raffensperger, K. and Shoykhet, M. (2021) Early Thalamic Injury After Resuscitation From Severe Asphyxial Cardiac Arrest in Developing Rats. Front. Cell Dev. Biol., 9. 10.3389/fcell.2021.737319

27. Shoykhet, M., Simons, D.J., Alexander, H., Hosler, C., Kochanek, P.M. and Clark, R.S.B. (2012) Thalamocortical dysfunction and thalamic injury after asphyxial cardiac arrest in developing rats. Journal of Neuroscience, 32. 10.1523/JNEUROSCI.5597-11.2012

28. Brenner, J.F., Dew, B.S., Horton, J.B., King, T., Neurath, P.W. and Selles, W.D. (1976) THE JOURNAL OF HISTOCHEMISTRY ANI) CYTOCHEMISTRY AN AUTOMATED MICROSCOPE FOR CYTOLOGIC RESEARCH A PRELIMINARY EVALUATION’.

29. Yagi, Y. (2011) Color standardization and optimization in Whole Slide Imaging. Diagn. Pathol., 6. 10.1186/1746-1596-6-S1-S15

30. Niazi, M.K.K., Parwani, A. V. and Gurcan, M.N. (2019) Digital pathology and artificial intelligence. Lancet Oncol., 20. 10.1016/S1470-2045(19)30154-8

31. Fernández-Arjona, M. del M., Grondona, J.M., Granados-Durán, P., Fernández-Llebrez, P. and López-Ávalos, M.D. (2017) Microglia morphological categorization in a rat model of neuroinflammation by hierarchical cluster and principal components analysis. Front. Cell. Neurosci., 11. 10.3389/fncel.2017.00235

